# Low heritability of crossover rate in wild sticklebacks

**DOI:** 10.1101/2022.05.26.493536

**Authors:** Mikko Kivikoski, Antoine Fraimout, Pasi Rastas, Ari Löytynoja, Juha Merilä

## Abstract

Crossover rate is mostly studied with domesticated or lab-reared populations and little is known about its genetic variation in the wild. We studied the variation and genetic underpinnings of crossover rate in outbred wild nine- (*Pungitius pungitius*) and three-spined (*Gasterosteus aculeatus*) sticklebacks. In both species, the crossover rate of females exceeded that of males as did also its repeatability (*R*_Females_ =0.21–0.33, *R*_Males_=0.026–0.11), implying individual differences of crossover rate in females, but no or less so in males. However, in both species and sexes additive genetic variance and heritability of crossover rate were effectively zero. A review of the previously reported repeatability and heritability estimates revealed that the repeatabilities in stickleback females were moderately high, whereas those in males were very low. Genome-wide association analyses recovered a few candidate regions possibly involved with control of crossover rate. The low additive genetic variance of crossover rate in wild sticklebacks suggest limited evolvability of crossover rate.

## Introduction

Populations’ ability to adapt to changing environmental conditions is tightly connected to the amount of genetic variability in traits influencing fitness (Lynch and Walsh 1998). While mutations are the ultimate source of genetic variation, crossovers recombine existing variation by breaking haplotypes and forming new ones. Hence, similarly to mutation rate (Lynch et al. 2016), crossover rate can be seen as a trait affecting evolvability (Zhong and Priest 2011, Charlesworth et al. 2017). Many previous studies have reported differences in crossover rate between sexes and individuals (Peñalba and Wolf 2020). Moreover, crossover rate has shown to be a heritable trait (Table 1), suggesting that it can evolve in response to natural selection. However, most of the available estimates are from domesticated or laboratory populations, and estimates from truly wild populations (i.e. populations not subject to any human interventions) are as yet lacking (Table 1).

**Table 1.**
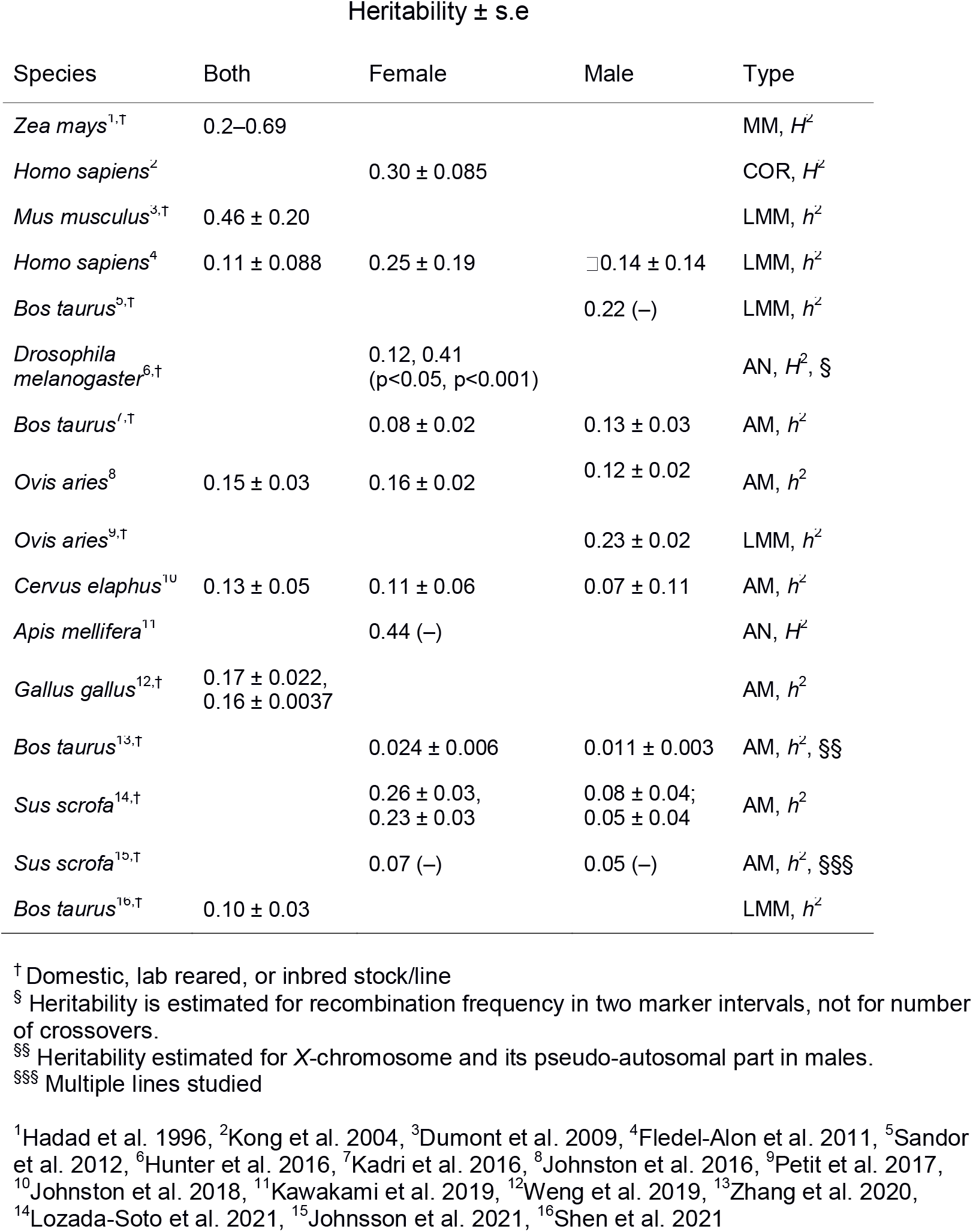
Synopsis of published estimates of heritability of crossover rate together with information on estimation method (*H*^2^ = broad-sense heritability, *h*^2^ = narrow-sense heritability, AM = animal model, AN = ANOVA, LMM = linear mixed model, MM = mid-parent mid-offspring regression, COR = sibling correlation). Empty cells indicate that the trait was not studied. For more details regarding sample size, see Table S1.

| Species | Heritability ± s.e | | | Type |
| --- | --- | --- | --- | --- |
|  | Both | Female | Male |  |
| *Zea mays*<sup>1,†</sup> | 0.2–0.69 | | | MM, $H^2$ |
| *Homo sapiens*<sup>2</sup> | | 0.30 ± 0.085 | | COR, $H^2$ |
| *Mus musculus*<sup>3,†</sup> | 0.46 ± 0.20 | | | LMM, $h^2$ |
| *Homo sapiens*<sup>4</sup> | 0.11 ± 0.088 | 0.25 ± 0.19 | □ 0.14 ± 0.14 | LMM, $h^2$ |
| *Bos taurus*<sup>5,†</sup> | | | 0.22 (–) | LMM, $h^2$ |
| *Drosophila melanogaster*<sup>6,†</sup> | | 0.12, 0.41<br>($p < 0.05$, $p < 0.001$) | | AN, $H^2$, § |
| *Bos taurus*<sup>7,†</sup> | | 0.08 ± 0.02 | 0.13 ± 0.03 | AM, $h^2$ |
| *Ovis aries*<sup>8</sup> | 0.15 ± 0.03 | 0.16 ± 0.02 | 0.12 ± 0.02 | AM, $h^2$ |
| *Ovis aries*<sup>9,†</sup> | | | 0.23 ± 0.02 | LMM, $h^2$ |
| *Cervus elaphus*<sup>10</sup> | 0.13 ± 0.05 | 0.11 ± 0.06 | 0.07 ± 0.11 | AM, $h^2$ |
| *Apis mellifera*<sup>11</sup> | | 0.44 (–) | | AN, $H^2$ |
| *Gallus gallus*<sup>12,†</sup> | 0.17 ± 0.022,<br>0.16 ± 0.0037 | | | AM, $h^2$ |
| *Bos taurus*<sup>13,†</sup> | | 0.024 ± 0.006 | 0.011 ± 0.003 | AM, $h^2$, §§ |
| *Sus scrofa*<sup>14,†</sup> | | 0.26 ± 0.03,<br>0.23 ± 0.03 | 0.08 ± 0.04;<br>0.05 ± 0.04 | AM, $h^2$ |
| *Sus scrofa*<sup>15,†</sup> | | 0.07 (–) | 0.05 (–) | AM, $h^2$, §§§ |
| *Bos taurus*<sup>16,†</sup> | 0.10 ± 0.03 | | | LMM, $h^2$ |

† Domestic, lab reared, or inbred stock/line
§ Heritability is estimated for recombination frequency in two marker intervals, not for number of crossovers.
§§ Heritability estimated for X-chromosome and its pseudo-autosomal part in males.
§§§ Multiple lines studied
<sup>1</sup>Hadad et al. 1996, <sup>2</sup>Kong et al. 2004, <sup>3</sup>Dumont et al. 2009, <sup>4</sup>Fledel-Alon et al. 2011, <sup>5</sup>Sandor et al. 2012, <sup>6</sup>Hunter et al. 2016, <sup>7</sup>Kadri et al. 2016, <sup>8</sup>Johnston et al. 2016, <sup>9</sup>Petit et al. 2017, <sup>10</sup>Johnston et al. 2018, <sup>11</sup>Kawakami et al. 2019, <sup>12</sup>Weng et al. 2019, <sup>13</sup>Zhang et al. 2020, <sup>14</sup>Lozada-Soto et al. 2021, <sup>15</sup>Johnsson et al. 2021, <sup>16</sup>Shen et al. 2021

Heritability is, by definition, a population specific measure (Visscher et al. 2008) and dependent on environmental conditions under which it is quantified (e.g. Falconer and Mackay 1996, Hoffmann and Merilä 1999). While heritabilities estimated under laboratory conditions may reflect poorly those in the wild (Riska et al. 1989, Weigensberg and Roff 1996), obtaining robust and unbiased estimates of crossover rate heritability from the wild is a challenging undertaking. First, it has been shown that both environmental conditions (e.g. Parsons 1988, Zhong and Priest 2011, Shen et al. 2021) and parental age can affect crossover rate (e.g. Martin et al. 2015), potentially influencing also estimates of its heritability. Second, as for any trait, choices in regard of experimental design and analytical methods may influence the estimates obtained (e.g. Falconer and Mackay 1996, Kearsey and Pooni 1996). In fact, the published heritability estimates of crossover rate have been obtained using quite diverse methods, for example with varying number of generations and family sizes (Table 1). Third, most estimates derive from domesticated populations which might have been subject to recent directional selection for higher crossover rate (Rees and Dale 1974, Burt and Bell 1987, Ross-Ibara 2004, but see Muñoz-Fuentes et al. 2015). Finally, crossover rate interpreted from genomic sequencing data (e.g. linkage maps) is a result of a long chain of analyses (e.g. read mapping, variant calling, allelic phasing), which can be sensitive to population demography, quality of the reference genome, sample size, number of informative markers and choice of the algorithms used (Rastas 2017).

The aims of this study were to estimate heritability and repeatability of crossover rate in the wild utilizing populations not subject to any kind of direct human interventions, as well as explore the genetic underpinnings of the observed variability with genome-wide association analyses. To this end, we took an advantage of the available high-density sex-specific linkage maps and high quality reference genomes (Kivikoski et al. 2021a) to obtain sex-specific estimates of repeatability (*R*) and narrow-sense (*h*^2^) heritability of the crossover rate for nine-(*Pungitius pungitius*) and three-spined (*Gasterosteus aculeatus*) sticklebacks. Both of these species have become important model systems in evolutionary biology, ecology and genetics (Bell and Foster 1994, Merilä 2013, Reid et al. 2021), including the study of the recombination process (Roesti et al. 2013, Rastas et al. 2016, Sardell et al. 2018, Shanfelter et al. 2019, Kivikoski et al. 2021a,b).

## Materials and methods

### Study populations, linkage maps and crossover data

The observed crossovers in both stickleback species were obtained from the linkage maps published earlier (linkage maps: Kivikoski et al. 2021a, offspring haplotypes: Kivikoski et al. 2021b). For the nine-spined stickleback, the parental fish were wild-caught individuals from the Baltic Sea, Helsinki (60°13’N, 25°11’E), Finland and they were artificially crossed in laboratory to produce the F_1_ fish (see: Kivikoski et al. 2021a and Rastas et al. 2016 for details of crossing and fish husbandry). Forty-one females were each crossed with two different males, and five females were each crossed with single (but different) males. For genotyping, parental fish were whole-genome sequenced with 5–10X coverage (Illumina Hiseq platforms, BGI Hong Kong). The genotyping of F_1_ fish was carried out with the DarTseq technology (Diversity Arrays Technology, Pty Ltd) as detailed in Rastas et al. (2016). Read mapping and variant calling were conducted with BWA-mem (ver. 0.7.15, Li 2013) and SAMtools mpileup (ver 1.9, Li et al. 2009) along the Lep-MAP3 software pipeline (Rastas 2017). The linkage maps were reconstructed from a data set of 133 parental fish forming 87 families with 938 F_1_-offspring in total. Linkage maps with 22,468 informative markers (i.e. markers informative to conclude crossover) were reconstructed with Lep-MAP3 software and number of paternal and maternal crossovers observed in F_1_-offspring were recorded from those, by counting the changes in haplotype phase. Four of the crosses had only a single offspring, and crossovers could not be inferred in those offspring. Therefore, those four single-offspring families were excluded from the analyses and the final dataset included 46 female parents (2–61 offspring per parent) and 83 male parents (2–38 offspring per parent), with 934 offspring in total.

For the three-spined stickleback, linkage maps were based on sequencing data available from Pritchard et al. (2017). Thirty males were each crossed with two different females generating 60 full-sib (both parents shared) and 30 half-sib (same father, different mothers) families in a North Carolina I type of breeding design (Kearsey and Pooni 1996). The number of offspring per parent was 8–20 and 3–10 for males and females, respectively. The wild caught parents originated from the Baltic Sea, Helsinki (60°13’N, 25°11’E), Finland and more details about the crossing scheme, rearing protocols and sequencing data are available from Pritchard et al. (2017) and Leder et al. (2015). Both parents and offspring were genotyped-by-sequencing following the Restriction-site Associated DNA (RAD) sequencing protocol of Elshire et al. (2011). The paired-end sequencing was carried out on the Illumina HiSeq2000 platform (see Pritchard et al. 2017 for details). Obtained RAD reads were mapped to the v4 reference genome of the three-spined stickleback (Peichel et al. 2017) using BWA-mem software, and the variants were called exactly as for the nine-spined stickleback along the Lep-MAP3 pipeline. In total, 28,187 informative markers were used for linkage map construction with Lep-MAP3 (see Kivikoski et al. 2021a for details).

For both species, the number of maternal and paternal crossovers were told from the changes in offsprings’ maternally and paternally inherited haplotypes, respectively, a change indicating crossover. The genomic positions of the crossovers (in base pairs) were defined as the midpoint of the informative markers indicating the crossover.

### Repeatability and sex-differences in crossover rate

We used univariate linear mixed models to study the repeatability and sex differences in crossover rate. Repeatability quantifies the proportion of phenotypic variance that is attributable to among individual variance (Nakagawa and Schielzeth 2010). In the model, the number of autosomal recombinations in the offspring was defined as a response variable, sex as fixed effect and parent identity as a random effect, so that recombinations observed in offspring were taken as an independent measurements of the parent’s crossover rate:

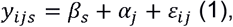

where *y*_*ijs*_ are the number of autosomal recombinations observed in offspring *i* of parent *j* of sex *s, β*_*s*_ is the fixed effect of the sex, *α*_*j*_ is the random effect of parent *j* and *ε*_*ij*_ is the random error. Random effects are assumed normally distributed, so that 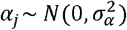, and 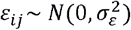. Repeatability, *R*, was calculated from the fitted model as 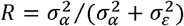, where 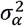 is the among individual variance (V_id_) and 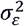 is the error variance (V_r_). In order to get sex-specific repeatability estimates, above model was fitted separately to male and female crossover data, with leaving out the sex as a fixed effect. For the model fitting, package MCMCglmm (ver. 2.32, Hadfield 2010) was used in R (ver. 4.1.1, R Core Team 2021). The model was run for 550,000 MCMC iterations, with a burn-in period of 50,000 iterations, and by sampling every 250th iteration. The model convergence was assessed by visual inspection of the trace plots and obtaining the effective sample sizes (R package CODA, ver. 0.19, Plummer et al. 2016) of each model component.

### Heritability of crossover rate

We estimated the overall and sex-specific heritability of crossover rate, i.e. the autosomal crossover count, using the ‘animal model’ approach (Kruuk 2004) as implemented in the MCMCglmm R package. Phenotypic variance (V_p_) in crossover rate was partitioned into additive genetic (V_a_), permanent environment (V_pe_) and residual components (V_r_) by fitting a model separately for both sexes in the form:

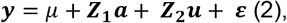

where ***y*** is the vector of phenotypic values (i.e. the number of autosomal recombinations in the offspring), *μ* is the population grand mean of the sex, ***z*_1_** and ***z*_2_** are the design matrices for additive genetic random effect and permanent environmental effect, respectively, ***a*** and ***u*** are the vectors of additive genetic and permanent environmental effects, respectively and ***ε*** is the vector of residual errors (e.g. Wilson et al. 2010). Narrow-sense heritability, *h*^2^, was calculated by dividing the additive genetic variance with the sum of all three variance components. For the overall heritability, sex was added as fixed effect, changing the equation to:

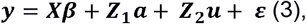

where ***X*** is the design matrix for the fixed effect, and ***β*** represents the means for the sexes. Elements of in *Z*_1_equations 2 and 3 were calculated from the SNP data by constructing the Genomic Relationship Matrix (GRM) among individuals. We used the method by VanRaden (2008) and built the reparameterized W matrix of genotyped markers such that:

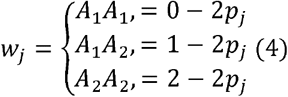

Where *j* is the vector of marker number and *p*_*j*_ is the frequency of the reference allele. The GRM was constructed as:

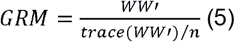

where *n* is the number of individuals, here linkage map parents. R package snpReady (ver. 0.9.6, Granato et al. 2018) was used to construct the GRM from autosomal loci. For the nine-spined stickleback, single nucleotide variants were joint-called as part of a larger set of nine-spined sticklebacks, covering populations from Europe, North-America and Russia. These data included 179 individuals from the Helsinki Baltic Sea population, including those 133 parental fish used in the linkage map reconstruction. From these joint-called data, the biallelic SNPs were kept. After this, the data were imputed and phased, using Beagle (ver. 4.1, Browning and Browning 2007, 2016). For the three-spined stickleback, the SNPs of the linkage map were used.

Prior to constructing the GRMs, we pruned the datasets from markers with a minimum allele frequency (MAF) lower than 0.01 and included only markers with a maximum of 10% missing data. This resulted in a total of 1,928,049 SNPs to be used in the nine-spined and 53,007 SNPs to be used in the three-spined stickleback analyses, respectively. Narrow-sense heritability was calculated as the ratio of additive genetic variance (V_a_) to total phenotypic variance (V_p_) for each sex and the precision of these estimates was retained by computing the 95% Highest Posterior Density (HPD) intervals using the HPDinterval function in MCMCglmm package.

For all heritability models, the number of MCMC iterations was set at 1,100,000, with a burnin period of 100,000 iterations. Every 500th iteration was used after thinning. The model convergence was assessed similarly to the repeatability models, by visual inspection of the trace plots and obtaining the effective sample size for the parameters.

### Genome-wide association analyses

A genome-wide association study (GWAS) was conducted to identify loci associated with crossover rate variation. As for the repeatability and heritability, the genome-wide association analyses were performed separately for both sexes and joined data. In the analysis of the joined data, sex was defined as fixed effect. The analysis was conducted with GenABEL (GenABEL project developers 2013, ver. 1.8) and RepeatABEL (Rönnegård et al. 2016, ver. 1.1) R packages on R version 3.4.4. GWAS was based on the same set of variants as the heritability estimation, with slightly different pruning criteria: only loci with minor allele count of five or more were used. This is equivalent of at least three individuals carrying the minor allele and the purpose was to avoid spurious signals. For the nine-spined stickleback, this left 2,227,185; 1,623,229 and 2,040,993 markers for combined, female and male data, respectively. For the three-spined stickleback, the corresponding numbers were 50,403; 41,854 and 33,999, respectively. The threshold of statistical significance was calculated by dividing the significance level (0.05) with the number of effective tests, estimated with program Keffective (Moskvina and Schmidt 2008). The number of effective tests were 1,346,555; 1,259,312 and 1,290,327 for the nine-spined stickleback and 43,615; 36,736; 29,322 for the three-spined stickleback. The nine-spined stickleback results were compared with the gene annotation by Varadharajan et al. (2019).

### Simulation assay

Observed recombinations are a sample of the crossovers that occurred in meiosis and may have different variance than the underlying trait, the number of crossovers in the bivalent (Weinstein 1936). Hence, the variance in the observed data must be studied with this in mind. We assessed whether the variance of the observed recombinations among the offspring could be a simple outcome of stochasticity in meiosis, even when the parents do not differ in their crossover rates in the bivalents. For this, we simulated data sets of maternal and paternal crossovers by assuming that all parents of the same sex have the same inferred average crossover frequency (see Fig. 2 in Kivikoski et al. 2021b). Assuming that all parents of same sex have identical crossover rate and that all offspring represent independent measurements of that trait, the data were simulated as follows: For 934 nine- and 517 three-spined stickleback offspring, the number of paternal or maternal crossovers in the bivalent was sampled from the chromosome and sex-specific multinomial distributions reported in Kivikoski et al. (2021b), which represent population average. After this, the number of observed crossovers was sampled from those crossover counts according to binomial factors (see Kivikoski 2021b for an example). For these generated data, the repeatability and variance of the total number of autosomal crossovers were estimated. Repeatability was estimated with the same model structure as for the empirical data, using frequentist approach implemented in lme4 R package (ver. 1.1, Bates et al. 2015), which allows quicker execution. The variance in the empirical data was compared to the distribution of 1,000 simulated variances.

### Synopsis of published heritability estimates of crossover rate

To put the estimated heritabilities of crossover rate into context, we conducted a literature review. This was done by searching articles reporting heritability of crossover rate (i.e. genome-wide recombination rate, autosomal crossover count) or recombination frequency (i.e. proportion of recombinant offspring) from Web of Science on 30 September 2021 with search terms ‘heritability AND recombination’ in the ‘Topic’ field, which directs search to title, abstract and author keywords. The search yielded 278 records which were refined to ‘Articles’, leaving 249 records. The search was repeated on 11 May 2022, but articles matching the criteria were not published after the initial search. The article titles and abstracts were manually searched (MK) for indication or direct report of the heritability estimates of crossover rate. Furthermore, in-text citations of articles reporting heritability estimates were searched for more reported literature references. Finally, articles that the authors of this manuscript were aware of by other means, were added. Condensed information of reported heritability estimates are listed in Table 1 and more detailed information in Table S1. Repeatability was not studied explicitly in all papers reporting heritability (e.g. Petit et al. 2017, Table S2), but it could be deducted from the animal model results by summing the proportion of total variance explained by additive genetic and permanent environmental effects (Wilson et al. 2010).

## Results

Differences in crossover rate between sexes were evident in both species and crossover rate varied among individuals of both species. The crossover rate was higher in females than males in both species (Fig. 1, Table 2). Furthermore, the trait was repeatable in females of both species and also in male three-spined sticklebacks, but not in male nine-spined sticklebacks (Table 2). Simulation assay confirmed that the repeatability estimates were not the result of inherent stochasticity, but reflect between-individual variation in the crossover rate (Tables S3 and S4). However, in male nine-spined sticklebacks the empirical repeatability was within the range yielded by the simulations, indicating that there are no (detectable) differences among individuals (Tables S3 and S4).

**Figure 1.**
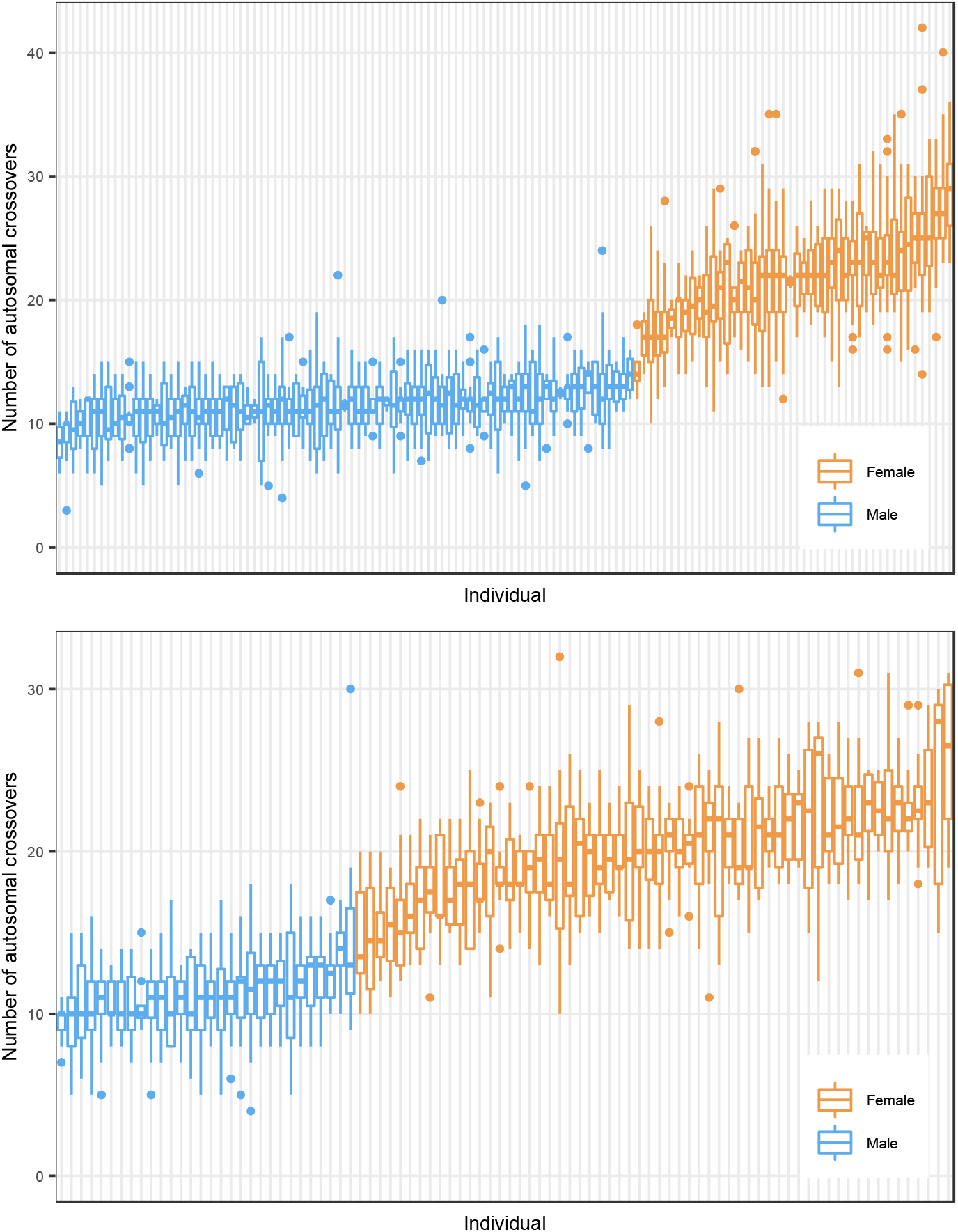
Number of observed autosomal crossovers in the nine-spined (top) and three-spined stickleback parents (bottom). Each data point is the number of crossovers detected in offspring that occurred in female (orange) and male (blue) parents. Thick horizontal line indicates mean and boxes and whiskers show the quartiles.

**Table 2.** Repeatability of crossover rate in male and female sticklebacks. Parameters (posterior mode) of fitted linear-mixed models with sex (‘analysis both’) as fixed effect and parental identity as random effect. Non-overlapping 95% credible intervals, i.e. highest posterior density intervals in parentheses in female and male intercepts as well as repeatability, confirm sex differences in repeatability of crossover rate. V_id_ = variance attributable to individual differences, V_r_ = residual variance component.

| Sex | Intercept female | Intercept male | $V_{id}$ | $V_r$ | Repeatability |
| --- | --- | --- | --- | --- | --- |
| *Pungitius pungitius* |  |  |  |  |  |
| Both | 21.87 (21.32–22.44) | 11.47 (11.05–11.89) | 2.56 (1.95–3.74) | 9.95 (9.34–10.62) | 0.22 (0.17–0.28) |
| Female | 21.84 (20.91–22.63) | – | 6.28 (4.09–11.12) | 13.20 (12.22–14.66) | 0.33 (0.24–0.46) |
| Male | – | 11.44 (11.25–11.64) | 0.16 (0.00–0.36) | 6.45 (5.79–6.98) | 0.026 (0.00–0.054) |
| *Gasterosteus aculeatus* |  |  |  |  |  |
| Both | 19.99 (19.44–20.41) | 11.20 (10.60–11.93) | 2.49 (1.72–3.95) | 9.42 (8.56–10.25) | 0.22 (0.16–0.30) |
| Female | 19.94 (19.31–20.50) | – | 3.35 (2.12–6.08) | 12.46 (11.11–14.44) | 0.21 (0.14–0.33) |
| Male | – | 11.24 (10.83–11.64) | 0.72 (0.25–1.57) | 6.19 (5.56–7.11) | 0.11 (0.036–0.20) |

**Table 3.** Heritability (*h*^*2*^) and variance component estimates (posterior modes) for female and male crossover rate in nine- and three-spined sticklebacks as estimated from animal models. V_a_ = additive genetic variance, V_pe_ = permanent environmental effect, V_r_ = residual variance, V_p_= phenotypic variance, n_FID_ =number of focal individuals (i.e. parents) and n_O_=number of offspring. 95% credible intervals are given in parentheses.

| Sex | $n_{FID}/n_o$ | $h^2$ | $V_a$ | $V_{pe}$ | $V_r$ | $V_p$ |
| --- | --- | --- | --- | --- | --- | --- |
| *Pungitius pungitius* |  |  |  |  |  |  |
| Both | 129/934 | 0.00061 (0.00–0.25) | 0.016 (0.00–3.25) | 0.013 (0.00–3.33) | 9.91 (9.40– 10.69) | 12.41 (11.75–13.92) |
| Female | 46/934 | 0.00069 (0.00–0.41) | 0.030 (0.00–9.38) | 0.031 (0.00–9.28) | 13.37 (12.22–14.73) | 20.57 (17.49–25.04) |
| Male | 83/934 | 0.00028 (0.00–0.039) | 0.0029 (0.00 –0.26) | 0.0022 (0.00 –0.26) | 6.28 (5.79–6.97) | 6.56 (5.96–7.14) |
| *Gasterosteus aculeatus* |  |  |  |  |  |  |
| Both | 90/517 | 0.00097 (0.00–0.26) | 0.017 (0.00–3.23) | 0.010 (0.00–3.33) | 9.18 (8.53–10.34) | 11.81 (10.81–13.56) |
| Female | 60/517 | 0.00085 (0.00–0.27) | 0.015 (0.00–4.61) | 0.024 (0.00–5.21) | 12.45 (11.18–14.28) | 16.28 (14.08–18.91) |
| Male | 30/517 | 0.00094 (0.00–0.16) | 0.0075 (0.00–1.22) | 0.0046 (0.00–1.27) | 6.28 (5.55–7.04) | 6.95 (6.26–8.20) |

The posterior probability distributions of heritability estimates were bimodal, except for nine-spined stickleback males (Fig. S1, S2). The posterior modes were close to zero, but in the three bimodal results another local maxima resided close to the repeatability estimates. In male nine-spined sticklebacks, the heritability was close to zero and the posterior distribution was unimodal (Fig. S1, S2).

Genome-wide association analyses of the nine-spined stickleback crossover rate revealed several suggestive peaks but the p-values were above the significance threshold for all peaks (Fig. 2). Corresponding analysis for the three-spined stickleback revealed one suggestive peak in chromosome 11, but due to low number of markers, the result had no statistical support (Fig. S5).

**Figure 2.**
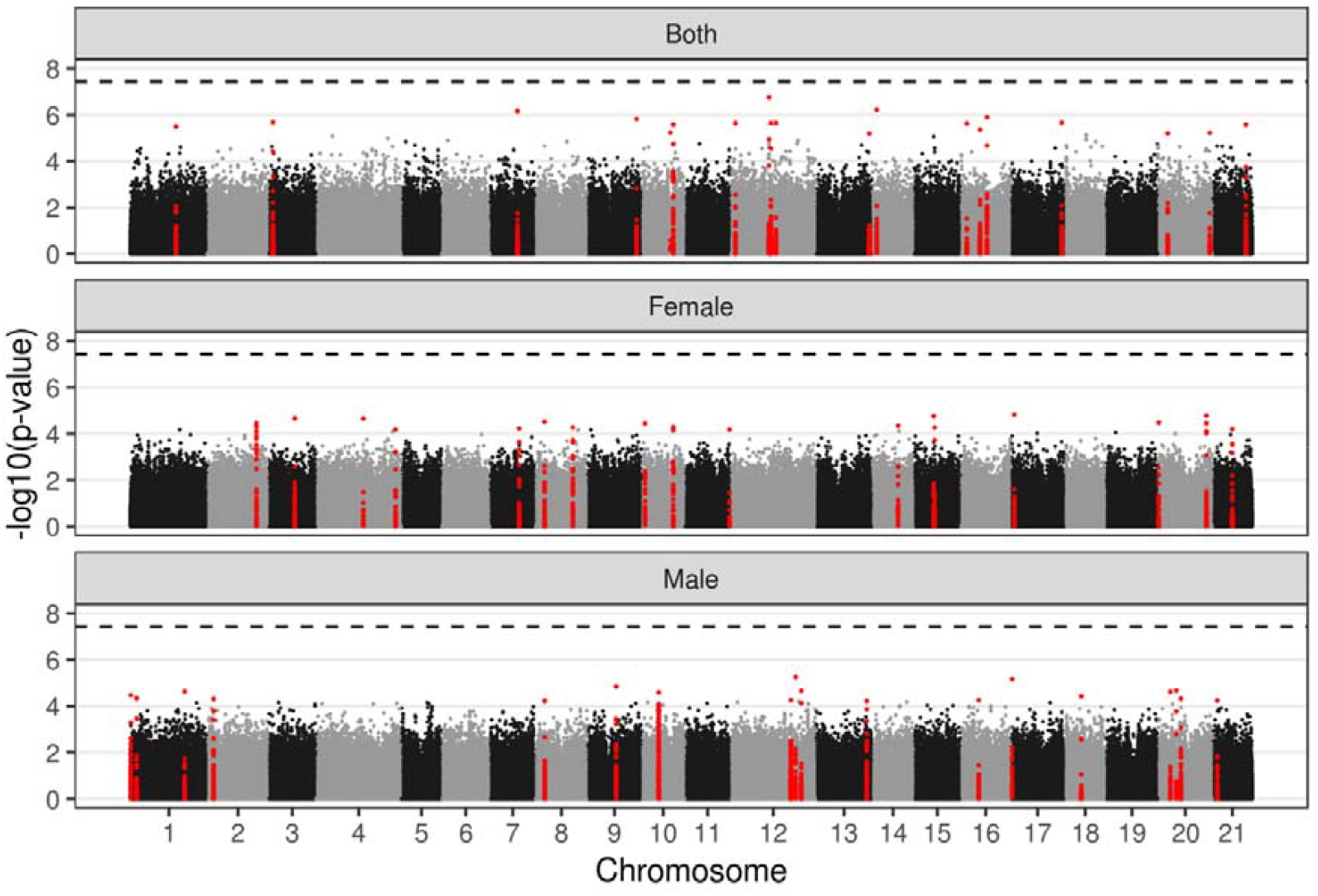
Results of genome wide association analysis of crossover rate in the nine-spined stickleback. The panels show p-values for each marker (n=1,598,255–1,679,069) when the trait was analyzed in both sexes (top), females (middle) and males (bottom). In each panel, 20 markers with the lowest p-value (highest on the plot) and the markers within the 5 kbp interval are colored red. Dashed line indicates the 5% significance level after correcting for the effective number of tests.

## Discussion

Assessing causes of variation in quantitative traits is a central task in evolutionary biology. One solution to this is decomposing phenotypic variance into different components, such as additive genetic and environmental effects, and estimating heritability. Heritability of crossover rate, namely the number of recombinations in the offspring, has been studied but mainly on livestock and laboratory models, whereas studies on wild populations are still scarce. With the aim to fill this knowledge gap, we assessed the repeatability and heritability of crossover rate in wild sticklebacks. The results revealed considerable among individual variation in female crossover rate and low among individual variation in male crossover rate, especially in the nine-spined stickleback. The repeatability estimates were within the range of the previous findings but at the extremes, especially in the nine-spined sticklebacks: reports of crossover rate repeatability exceeding 0.30 are rare, as are those close to zero (Table S2). We also found marked sexual dimorphism in repeatability of crossover rate, something that has been seldom observed in previous studies (Table S2).

Although there were individual differences in crossover rate in females of both species and in male three-spined sticklebacks, those differences seemed not to be underpinned by much additive genetic variance. Lack of (estimable) additive genetic variance led to bimodal posterior distributions of the female and overall heritability, indicating that the MCMCglmm algorithm did not converge into solution when decomposing the among individual variance into additive genetic (*V*_*a*_) and environmental (*V*_*pe*_) components. More specifically, for the female parents with crossover rate deviating from the grand mean, the posterior distributions of the additive genetic and permanent environment random effects were bimodal, with one local maxima close to zero (i.e. grand mean, as expected when there is no additive genetic variation) and another one at the difference between the parents’ mean and the grand means (as suggested when breeding values cause the difference). In line with this, the estimates of the two random effects (*V*_*a*_ and *V*_*pe*_) were negatively correlated over the MCMC iterations (Fig. S4). In other words: the variance that was not attributable to the residual variance component, *V*_*r*_, fluctuated between *V*_*a*_ and *V*_*pe*_, depending on the iteration. Despite the bimodality of *V*_*a*_ and *V*_*pe*_ estimates, the posterior distribution of the residual variance, *V*_*r*_, was unimodal. Furthermore, the posterior modes of *V*_*r*_ were effectively identical in repeatability and heritability analyses. The consistency of *V*_*r*_ estimates in the two analyses and the simulation assay confirm that the repeatability estimates are robust and indicate that there is consistent individual variation in the crossover rate among females of both species and male three-spined sticklebacks. As the repeatability sets the upper-limit for heritability (Kruuk 2004, but see Dohm 2002), these results indicate that the heritability in female sticklebacks is at best moderate, whereas that of males is very low.

Here, as in many recent studies (e.g. Zhang et al. 2020, Lozada-Soto et al. 2021), the phenotype was the *number of recombinations in the offspring*, not the number of crossovers in meiosis. As can be gleaned from Tables 1 and S2, heritability and repeatability estimates of this trait are commonly below 0.3 indicating that most of the phenotypic variance in the trait goes unexplained. We argue that this is largely due to the stochastic association between the number of crossovers in meiosis and number of recombinations in offspring. As only two chromatids are involved in one crossover, the number of recombinations in the offspring is a sample of crossovers that occurred in the meiosis (Weinstein 1936). Therefore, gametes originating from meioses with the same number of crossovers will carry a variable number of recombinations just by chance. Furthermore, the number of crossovers in the bivalent varies among meioses (see Kivikoski et al. 2021b for example), which further increases the variance in the number of recombinations in offspring. Multiple measurements from the same individual vary inevitably, for example due to measurement error (e.g. Merilä and Björklund 1995) or changes in environment. However, here the inherent stochasticity causes phenotypic variance within the individual itself even in absence of other such factors, which makes the upper limit for repeatability and heritability of crossover rate less than one. Therefore, the ‘unexplained’ variance in the phenotype must be interpreted with caution. This point has been seldom addressed explicitly, but it was made by Valentin (1973) for recombination fractions.

Genome wide association analyses aiming to map genomic regions potentially involved in control of crossover rate variation revealed several suggestive regions in the nine-spined stickleback, but none of those reached the significance thresholds likely due to low sample size. Both females and males had one suggestive candidate in LG10, close to the *Top2b* gene, which has been reported in a similar association analysis on red deer (Johnston et al. 2018). However, the loci yielding the association were on different sides of the gene’s start position in males and females. For females and when sexes were analyzed together, another candidate region was found in LG21 close to *Rec8* gene, whose association with recombination has been reported in cattle (Sandor et al. 2012, but see Kadri et al. 2016). For males, one candidate loci was in LG9 close to *Prdm9* gene, which is known to affect crossover hotspots (e.g. Baudat et al. 2010), and that has also been reported in a similar study on bulls (Sandor et al. 2012). In total, the genome wide association analysis suggests that genes that are previously found to be associated with recombination rate in vertebrates, may be associated in the wild sticklebacks as well. For female nine-spined sticklebacks, one suggestive peak was found in LG2, but none of the genes reported in earlier studies was located in that chromosome. In the three-spined sticklebacks the analysis did not reveal clear signals, likely due to smaller sample size and fewer SNPs.

Despite recent efforts to study repeatability and heritability of crossover rate, there is need for further research. For example, studies from the wild are scarce and the stochastic nature of crossover data has obtained very little attention. Although the recombinations here were observed in lab reared offspring, the parents in which the crossovers occurred had lived in the wild and the data should reflect phenotype in natural habitat. Furthermore, family sizes of the studied species are very different (Table S1), which affects the power of animal models with repeated measurements. The among individual variance in stickleback females, without any detectable additive genetic contribution indicates that the individual differences are largely attributable to environmental or non additive genetic effects. This does not imply total lack of standing genetic variation in (female) recombination rate: it could be very low or organized in the way that the breeding values of the individuals are effectively the same, but our data is probably too small to get an accurate estimate of low heritability. However, nine-spined stickleback males had no individual differences. Hence, it is possible that female crossover rates could evolve more readily in response to selection than those of males. This is intriguing in the perspective that traits closely associated with fitness tend to have lower heritabilities than traits less closely associated with fitness (e.g. Merilä and Sheldon 2000). It is tempting to speculate that low repeatability and heritability (and V_a_) in the males could be indicative of strong selection to maintain low recombination in males. As the males have the obligate crossover in meiosis (Kivikoski et al. 2021b), the lower boundary for the crossover rate could be set by the proper segregation of chromosomes (Lenormand et al. 2016). Hence, the low heritability and phenotypic variance in male crossovers suggests that some selective agent suppresses the male crossover rate close to the bare minimum.

## Conclusions and future directions

To sum up, the results show that crossover rate varies marginally among male but more so among female sticklebacks. However, the heritability of crossover rate was at best low in both females and males. Low heritability does not imply total lack of standing genetic variation underlying crossover rate, which is also indicated by results of the genome wide association analyses: genes reported to be associated with recombination rate variation in other organisms appeared to be associated (albeit non significantly) with the crossover rate in the sticklebacks as well. Potentially interesting lines of future research would involve estimation of the heritability of crossover rate from data spanning multiple generations and in small isolated pond populations of nine-spined stickleback in particular. This because populations - and in particularly small ones (Otto and Barton 2001) - adapting to new environments can be predicted to benefit from increased recombination rate generating novel combinations of traits (Burt and Bell 1987), but also to allow selection to more easily dissociate and purge harmful mutations segregating in these small and inbred populations.

## Acknowledgements

We thank all the people whose contributions to fish rearing and laboratory work made this study possible. The authors would also like to thank Frédéric Guillaume for commenting the manuscript and Marion Sinclair-Waters and Petri Kemppainen for advice on GWAS. Jon Slate is thanked for suggesting a repeatability analysis. Our research was supported by Academy of Finland (grants: 129662, 134728 and 218343 to JM; 322681 to AL; 343656 to PR), the Doctoral Programme in Wildlife Biology Research (University of Helsinki, funding to MK), Alfred Kordelin Foundation (grant 210190 to MK) as well support from the Helsinki Institute of Life Science (HiLife, grant to JM). We wish to acknowledge CSC – IT Center for Science, Finland, for access to computational resources.

## Data accessibility statement

Genomic sequencing data for the nine-spined stickleback parents and offspring have been published earlier (Kivikoski et al. 2021a) and they are available in European Nucleotide Archive (ENA) under the accession numbers PRJEB39736 and PRJEB39760, respectively. The sequencing reads for the three-spined stickleback have been published earlier (Pritchard et al. 2017) and they are available in NCBI Sequence Read Archive with BioProject ID PRJNA340327. The computer code and linkage maps with the offspring haplotypes are available in GitHub: github.com/mikkokivikoski/recombinationStudies.

## Author contributions

MK led the writing of the manuscript, designed the simulation assay, and executed the analyses. MK and AF designed the animal model and the relatedness matrix construction. PR made the linkage map and AL performed the variant calling for the larger set of nine-spined sticklebacks. The study was supervised by JM and AL. All authors participated the manuscript writing.

