## Supplementary material for "Low heritability of crossover rate in wild sticklebacks"

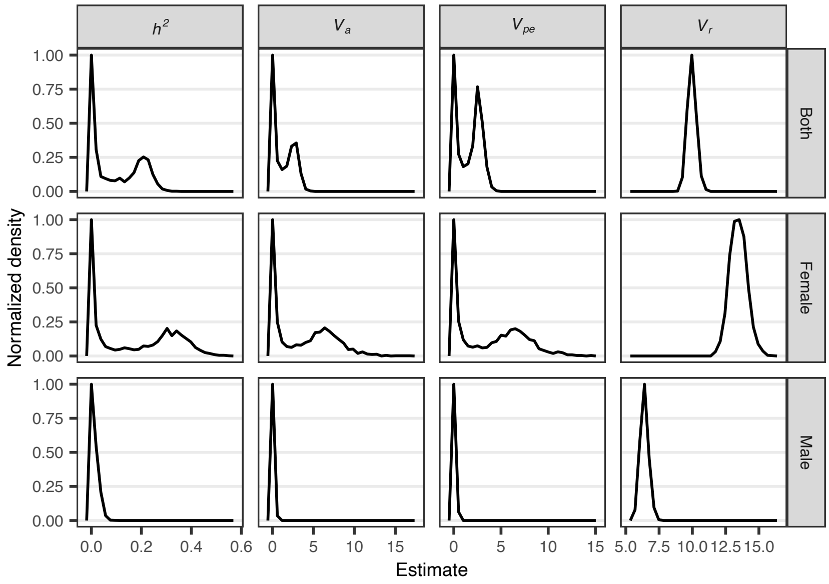


**Fig S1.** Posterior probability density functions of crossover rate heritability ($h^2$) and variance components for the nine-spined stickleback (*Pungitius pungitius*). $h^2$= narrow-sense heritability, $V_a$ = additive genetic variance, $V_{pe}$ = permanent environmental variance, $V_r$ = residual variance.

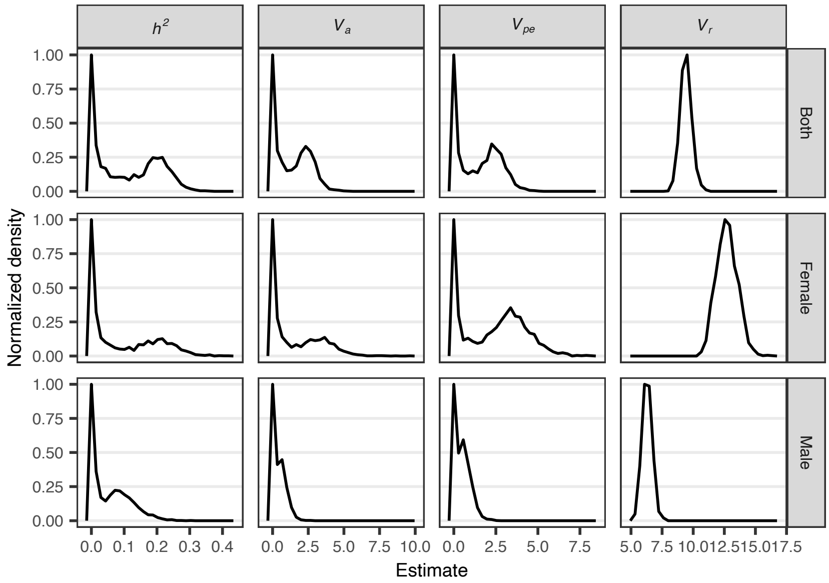


**Fig S2.** Posterior probability density functions of crossover rate heritability ($h^2$) and variance components for female and male three-spined sticklebacks (*Gasterosteus aculeatus*). $h^2$= narrow-sense heritability, $V_a$ = additive genetic variance, $V_{pe}$ = permanent environmental variance, $V_r$ = residual variance.

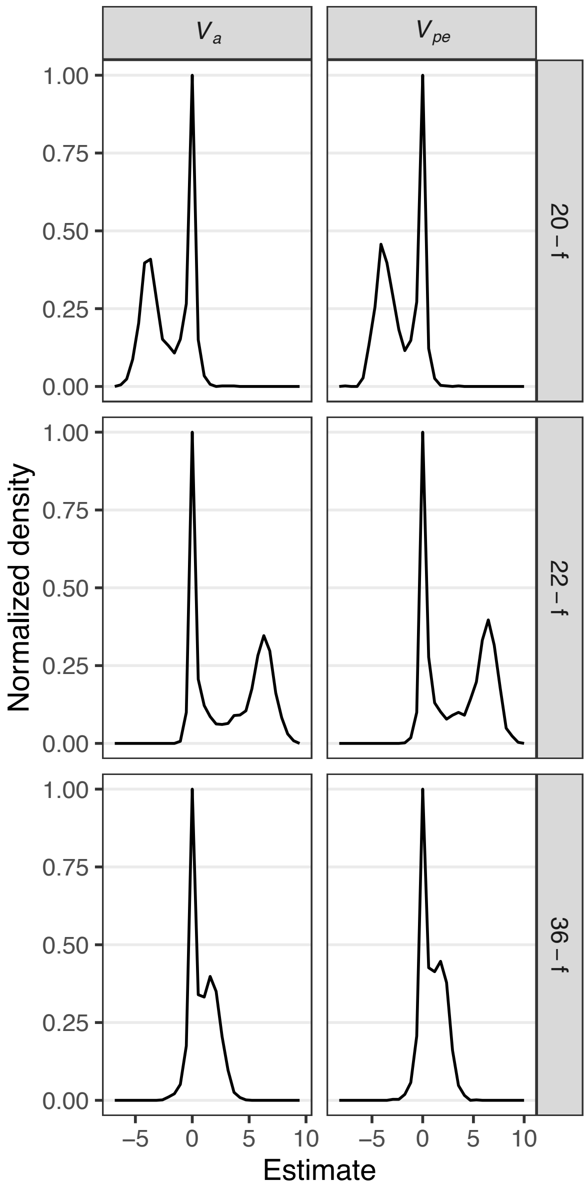


**Fig S3.** Three examples of posterior probability density functions of additive genetic ($V_a$) and permanent environment ($V_{pe}$) random effects of nine-spined stickleback females.

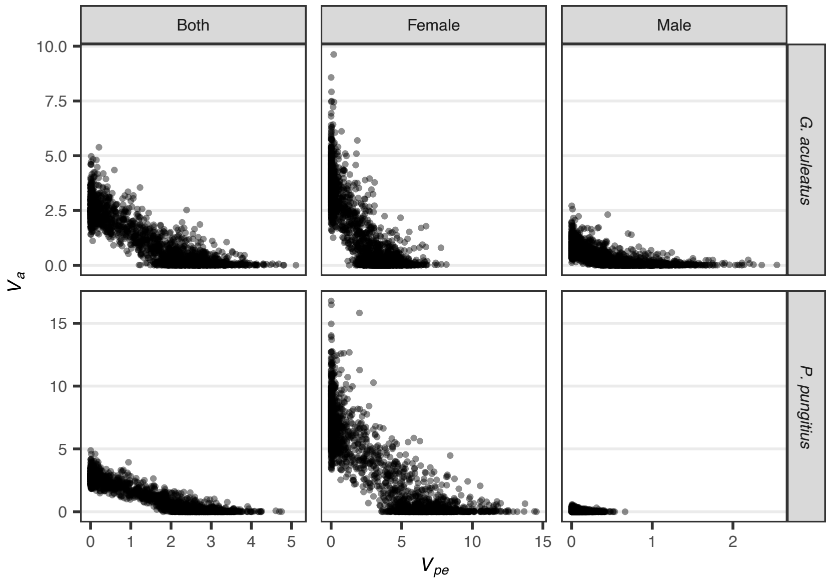


**Fig. S4** Negative correlation of additive genetic variance ($V_a$) and permanent environmental ($V_{pe}$) variance components in the two stickleback species. Each dot (n=2,000) represents an estimate of one iteration of the Markov chain Monte Carlo of the animal model used for heritability estimation.

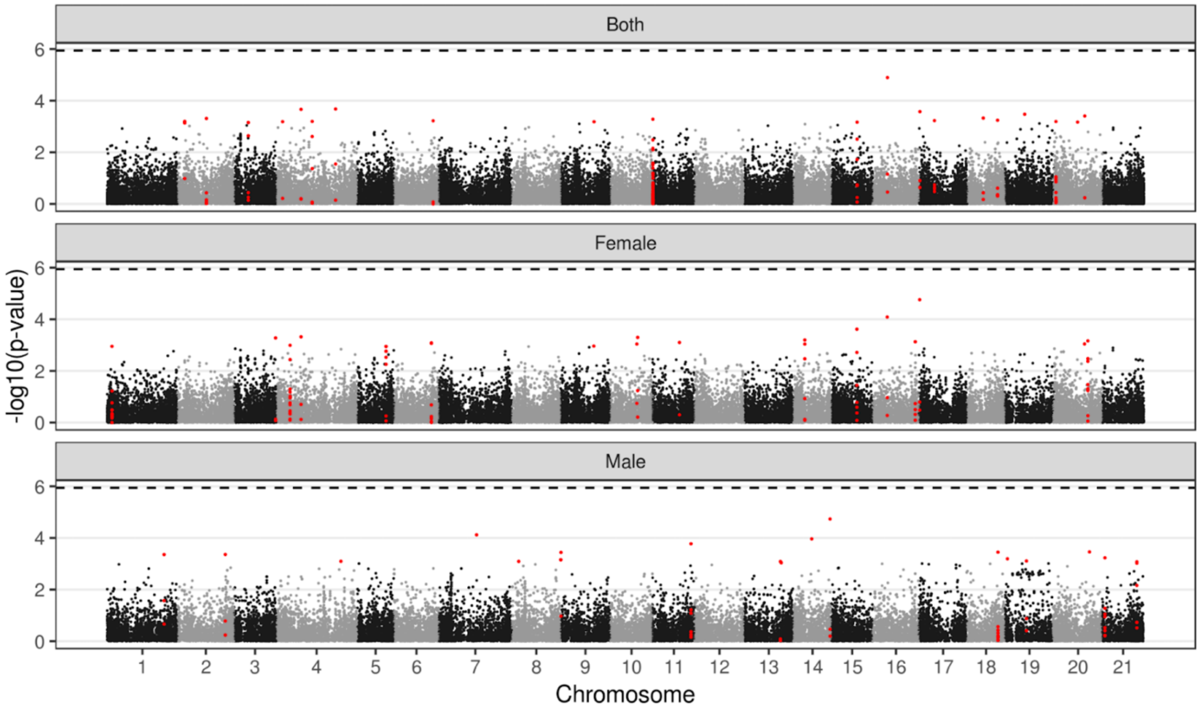


**Fig. S5** Results of genome-wide-association analyses of crossover rate in the three-spined stickleback. The panels show p-values for each marker (n=1,598,255–1,679,069) when the trait was analyzed in both sexes (top), females (middle) and males (bottom). In each panel, 20 markers with the lowest p-value (highest on the plot) and the markers within the 5 kbp interval are colored red. Dashed line indicates the significance level after correcting with the effective number of tests.

**Table S1. Sample sizes in the recombination rate heritability studies (Table 1 in the main text).** n refers to sample size, $n_{p\text{-}o}$=number of pairs in parent-offspring regression, $n_f$=number of families,$n_l$=number of lines, $n_c$=number of colonies, $n_d$=number of drones, $n_{fid}$=number of focal individuals in animal model, $n_o$=number of offspring

| Species | Both | Female | Male |
|---|---|---|---|
| *Zea mays*[1] | $n_{p\text{-}o}$=6 | | |
| *Homo sapiens*[2] | | $n_{pairs}$ = 554 | |
| *Mus musculus*[3] | $n_f$=8, $n_o$=2,474 | | |
| *Homo sapiens*[4] | $n_f$=163, $n_o$=ca. 753 | $n_f$=163, $n_o$=ca. 753 | $n_f$=163, $n_o$=ca. 753 |
| *Bos taurus*[5] | | | $n_p$=762, $n_o$=10,106 |
| *Drosophila melanogaster*[6] | | $n_l$=112<br>3 replicates per line | |
| *Bos taurus*[7] | | $n_f$=3,386 | $n_f$=2,135 |
| *Ovis aries*[8] | $n_{fid}$=813, $n_o$=3,330 | $n_{fid}$=586, $n_o$=2,134 | $n_{fid}$=227, $n_o$=1,196 |
| *Ovis aries*[9] | | | n=354 |
| *Cervus elaphus*[10] | $n_{fid}$=337, $n_o$=1,341 | $n_{fid}$=256, $n_o$=859 | $n_{fid}$=81, $n_o$=482 |
| *Apis mellifera*[11] | | $N_c$=16, $n_d$=158 | |
| *Gallus gallus*[12] | | $n_{fid}$=166, $n_o$=332;<br>$n_{fid}$=1096, $n_o$=4,573 | $n_{fid}$=282, $n_o$=969;<br>$n_{fid}$=621, $n_o$=4,719 |
| *Bos taurus*[13] | | $n_p$=10,737, $n_o$=114,254 | $n_p$=2,839, $n_o$=114,254 |
| *Sus scrofa*[14] | | $n_{fid}$=1,755, $n_o$=5,706;<br>$n_{fid}$=1,356, $n_o$=4,628 | $n_{fid}$=270, $n_o$=5,718;<br>$n_{fid}$=281, $n_o$=4,657 |
| *Sus scrofa*[15] | | $n_{fid}$=727–5,171 | $n_{fid}$=78–492 |
| *Bos taurus*[16] | $n_f$=36,009 | | |

For the references, see the legend of Table 1 in the main text.

**Table S2.** Synopsis of published estimates of repeatability (s.e.) of crossover rate from studies using animal model.

| Species | Overall | Female | Male |
|---|---|---|---|
| *Bos taurus*[7] | – | 0.09 | 0.18 |
| *Ovis aries*[8,§] | 0.15 (0.02) | 0.16 (0.02) | 0.12 (0.03) |
| *Cervus elaphus*[10,§] | 0.18 (0.03) | 0.16 (0.04) | 0.13 (0.05) |
| *Apis mellifera*[11] | – | 0.44 (–) | – |
| *Gallus gallus*[12] | 0.21, 0.24 (0.02, 0.03) | – | – |
| *Bos taurus*[13] | – | 0.09 (0.01) | 0.18 (0.01) |
| *Sus scrofa*[14] | – | 0.42, 0.39 (0.02, 0.02) | 0.27, 0.28 (0.03, 0.03) |
| *Sus scrofa*[15,§] | – | <0.15 | <0.15 |

[§]Repeatability was not reported explicitly in the study. Here, inferred as 1 – error variance. For the references, see the legend of Table 1 in the main text.

**Table S3** Statistics of crossover count data from 934 nine-spined stickleback offspring of 129 parents (46 females and 83 males). 95% quantiles for the corresponding values in 1,000 simulated data sets in parentheses.

| Statistic | Female (n=46) | Male (n=83) |
|---|---|---|
| Population grand mean (n=934) | 22.02 (21.56–22.02)§ | 11.46 (10.97–11.28)§ |
| Population variance (n=934) | 20.96 (11.70–14.01)§ | 6.53 (5.58– 6.71)§ |
| Variance of parental means | 8.09 (0.68–2.21)§ (1.10–3.68)§§ | 0.88 (0.63–1.37)§ (0.69–1.47)§§ |
| Repeatability | 0.33 (0.00– 0.018)§ (0.00–0.025)§§ | 0.022 (0–0.027)§ (0.00–0.025)§§ |

§Derived from simulated data assuming the same crossover rate in all parents.
§§Derived from randomized data.

**Table S4** Statistics of crossover count data from 517 three-spined stickleback offspring of 90 parents (60 female and 30 male). 95% quantiles for the corresponding values in 1,000 simulated data sets in parentheses.

| Statistic | Female (n=60) | Male (n=30) |
|---|---|---|
| Population grand mean (n=517) | 19.86 (19.46–20.04)§ | 11.21 (10.81–11.25)§ |
| Population variance (n=517) | 16.25 (9.98–12.80)§ | 7.04 (5.41–6.88)§ |
| Variance of parent means | 5.65 (0.96–2.01)§ (1.00–2.65)§§ | 1.19 (0.20–0.60)§ (0.22–0.63)§§ |
| Repeatability | 0.21 (0.00–0.046)§ (0.00–0.048)§§ | 0.11 (0–0.034)§ (0.00–0.031)§§ |

§Derived from simulated data assuming the same crossover rate in all parents.
§§Derived from randomized data.
